# HIT-scISOseq: High-throughput and High-accuracy Single-cell Full-length Isoform Sequencing for Corneal Epithelium

**DOI:** 10.1101/2020.07.27.222349

**Authors:** Ying-Feng Zheng, Zhi-Chao Chen, Zhuo-Xing Shi, Kun-Hua Hu, Jia-Yong Zhong, Chun-Xiao Wang, Wen Shi, Ying Chen, Shang-Qian Xie, Feng Luo, Xiao-Chen Bo, Chong Tang, Yi-Zhi Liu, Chuan-Le Xiao

## Abstract

Single-cell isoform sequencing can reveal transcriptomic dynamics in individual cells invisible to bulk- and single-cell RNA analysis based on short-read sequencing. However, current long-read single-cell sequencing technologies have been limited by low throughput and high error rate. Here we introduce HIT-scISOseq for high-throughput single-cell isoform sequencing. This method was made possible by full-length cDNA capture using biotinylated PCR primers, and by our novel library preparation procedure that combines head-to-tail concatemeric full-length cDNAs into a long SMRTbell insert for high-accuracy PacBio sequencing. HIT-scISOseq yields > 10 million high-accuracy full-length isoforms in a single PacBio Sequel II 8M SMRT Cell, providing > 8 times more data output than the standard single-cell isoform PacBio sequencing protocol. We exemplified HIT-scISOseq by first studying transcriptome profiles of 4,000 normal and 8,000 injured corneal epitheliums from cynomolgus monkeys. We constructed dynamic transcriptome landscapes of known and rare cell types, revealed novel isoforms, and identified injury-related splicing and switching events that are previously not accessible with low throughput isoform sequencing. HIT-scISOseq represents a high-throughput, cost-effective, and technically simple method to accelerate the burgeoning field of long-read single-cell transcriptomics.

## Introduction

Single-cell RNA sequencing (scRNA-Seq) technologies have emerged as a powerful tool for resolving cellular heterogeneity in developmental biology, oncology, neuroscience, immunology, and many fields that involve complex biological and pathological processes^1–5^. The current mainstream scRNA-seq technologies rely on next-generation sequencing (NGS) platform ^6^. While accurate and cost-effective, these short-read technologies are mainly used for quantitative analysis at the single-cell gene level, and they cannot resolve complex transcript isoforms and characterize alternative splicing, chimeric transcript, and sequence diversity^7^. Shortly after NGS, third-generation sequencing (TGS) technologies have emerged with their distinctive feature of long single-molecule sequencing reads, offering some advantages over short-read sequencing^8^. Building upon TGS, a single-cell full-length isoform sequencing method (scISO-seq) is being gradually incorporated in the research setting^9^. This method is made possible with the use of droplet-based systems such as 10x Genomics to generate full-length cDNAs, which are then proceeded to yield full-length transcriptome sequencing data using TGS platforms (Pacific Biosciences (PacBio) or Oxford Nanopore Technologies (ONT))^9–11^. scISO-seq enables the identification and quantification of full-length mRNAs without the need for assembly and imputation. However, it typically suffers from low yields and high costs, posing a limit on the accessibility and scalability of the technology.

There are some technical hurdles in improving the throughput of scISO-seq. For the Nanopore PromethION platform, while the throughput of the system is significantly increased (>20 million raw reads), the error rate of reads is still around 15%, complicating the cell-barcode demultiplex efficiency (14% - 40%)^12^. With 7 PromethION flow cells, one could only characterize < 1,000 single cells, and the data still require Illumina-based error correction^10, 13^. For the PacBio platform, the raw subreads error rate was equally high. To increase sequencing accuracy, PacBio has recently released a circular consensus sequencing (CCS) system that could achieve high accuracy (false positive rate of cell barcode <0.1%)^14^. However, the throughput of PacBio CCS system remains low because of the following issues: Firstly, in the 10x Genomics single-cell preparation pipeline, a high proportion (around 50%) of undesirable cell-barcode-free reads (TSO artifacts) would be introduced during library construction. Having these artifacts sequenced would result in a 50% waste in TGS sequencing resource^15^. Secondly, in a conventional CCS SMRTbell library preparation procedure, one would obtain a short-insert (around 2Kb) cDNA library, as it is typically the average length of a human transcript. The short inserts, however, are misaligned with PacBio’s HiFi long-read sequencing capacity (9-15 Kb per CCS)^14^, thus impairing Sequel II system’s sequencing outputs. It is therefore not surprising that, in a previous study based on PacBio Sequel sequencing platform, a total of 11 Sequel 1M SMRT cells were needed to generate 5.2 million reads to characterize 6,000 single cells^16^.

To overcome these limitations and unlock the full potential of CCS technology, we describe high-throughput single-cell isoform sequencing (HIT-scISOseq), a strategy for high-throughput and high accuracy single-cell isoform sequencing. This novel method is made possible by the following steps. Firstly, we used a PCR-based biotin-assisted capture procedure for removing TSO artifacts and enrichment of single-cell full-length cDNAs sequences. We showed that this could effectively reduce the proportion of TSO artifacts to less than 8%. Secondly, we proposed a novel library preparation procedure, with a head-to-tail concatenation of multiple full-length cDNAs into one long SMRTbell insert for Sequel II sequencing. The extended-length circular template maximizes the yield in each nanoscale observation chamber (Zero Mode Waveguide, ZMW), significantly increasing the consensus reads for full-length isoform output. Finally, we developed a computational pipeline to enable rapid and accurate analysis of scISOseq data generated from HIT-scISOseq, which poses a computational challenge for mapping the linked transcript reads. Using HIT-scISOseq, we generated > 10 million full-length cDNAs with one single Sequel II SMRT cell, 8times of the yields from a standard scISOseq approach. We analyzed transcriptomes from thousands of corneal epithelial cells, and identified cell-type-specific expression and known and novel isoforms. HIT-scISOseq is accurate, inexpensive and technically simple, making it accessible for a wide range of large-scale high-throughput transcriptome projects.

## Results

### Design principle of HIT-scISOseq

Droplet-based single-cell RNA sequencing, especially the 10x Chromium system, is increasingly used as a scalable solution for full-length cDNA library construction in scISOseq, through the application of microfluidic partitioning to capture mRNA in single cells and to prepare barcoded full-length cDNA libraries for PacBio sequencing platform (Figure S1). This approach has enabled a detailed look at the facets of single-cell transcriptomes from hundreds to tens of thousands of cells that have been previously out of reach in Illumina short-read sequencing technologies. However, some well-known by-products may occur by combining the template-switching oligonucleotides (TSOs) and reverse transcription reactions during the preparation of small volume cDNA libraries. The combined procedure could yield 40-50% of barcode-less TSO artifact libraries, consequently accounting for 50% of artifact reads. Another significant barrier to high-throughput CCS read yield is the short insert size of SMRTbell library on Sequel II systems. Recent advance now enables 10-15 kb SMRTbell libraries for HiFi Long Read Sequencing, in which a single DNA polymerase enzyme is affixed at the bottom of a ZMW nanoscale well with a single molecule of DNA as a template. For genome sequencing, PacBio recommends a library insert size of 9-15 kb for the Sequel II system. However, methods of preparing cDNA transcripts have been limited by the short cDNA insert size (on average 3 kb), which is not compatible with the long-read capacity of ZMW nanoscale well. Currently, the productivity of scISOseq has been 20%-30% of those from gDNA sequencing. The low throughput increases cost and hampers the application of full-length sequencing for accurate transcript identification at the isoform level.

To enable scalable sequencing of human transcriptomes, we describe HIT-scISOseq (Figure 1), a high-throughput full-length isoform sequencing method that consists of two phases: a capture phase and a concatenation phase. First, in order to remove the impact of TSO artifacts, we constructed a biotinylated PCR primer that hybridizes to the desirable cDNAs, which could then be biotinylated during the PCR amplification and then captured with Streptavidin beads (Figure 1B-C). Second, in order to create a long-insert SMRTbell library compatible with the capacity of Sequel II ZMW nanoscale well, we extended the length of SMRTbell template by linking multiple cDNA inserts and joined them together in a head-to-tail manner for downstream Sequel II CCS sequencing. The linking was made possible by designing a palindrome sequence at the 5’ end of the primer in the second round of PCR, and by the use of USER enzyme to generate sticky ends (Figure 1D). Multiple cDNAs were joined using DNA ligase in a head to tail fashion (Figure 1E). With our protocol, HIT-scISOseq could be performed in a widely accessible Drop-seq chromium system by 10x Genomics, and provides essential isoform information with a significant reduction of TSO artifacts, from 50% in standard scISOseq method to 8% in HIT-scISOseq (Figure 1H), and with a marked increase in mapped full-length reads to 8 times of that associated with standard scISOseq method (Figure 1I). We demonstrate that this approach is reproducible and readily adaptable to high-throughput single-cell isoform sequencing applications.

**Figure 1.**
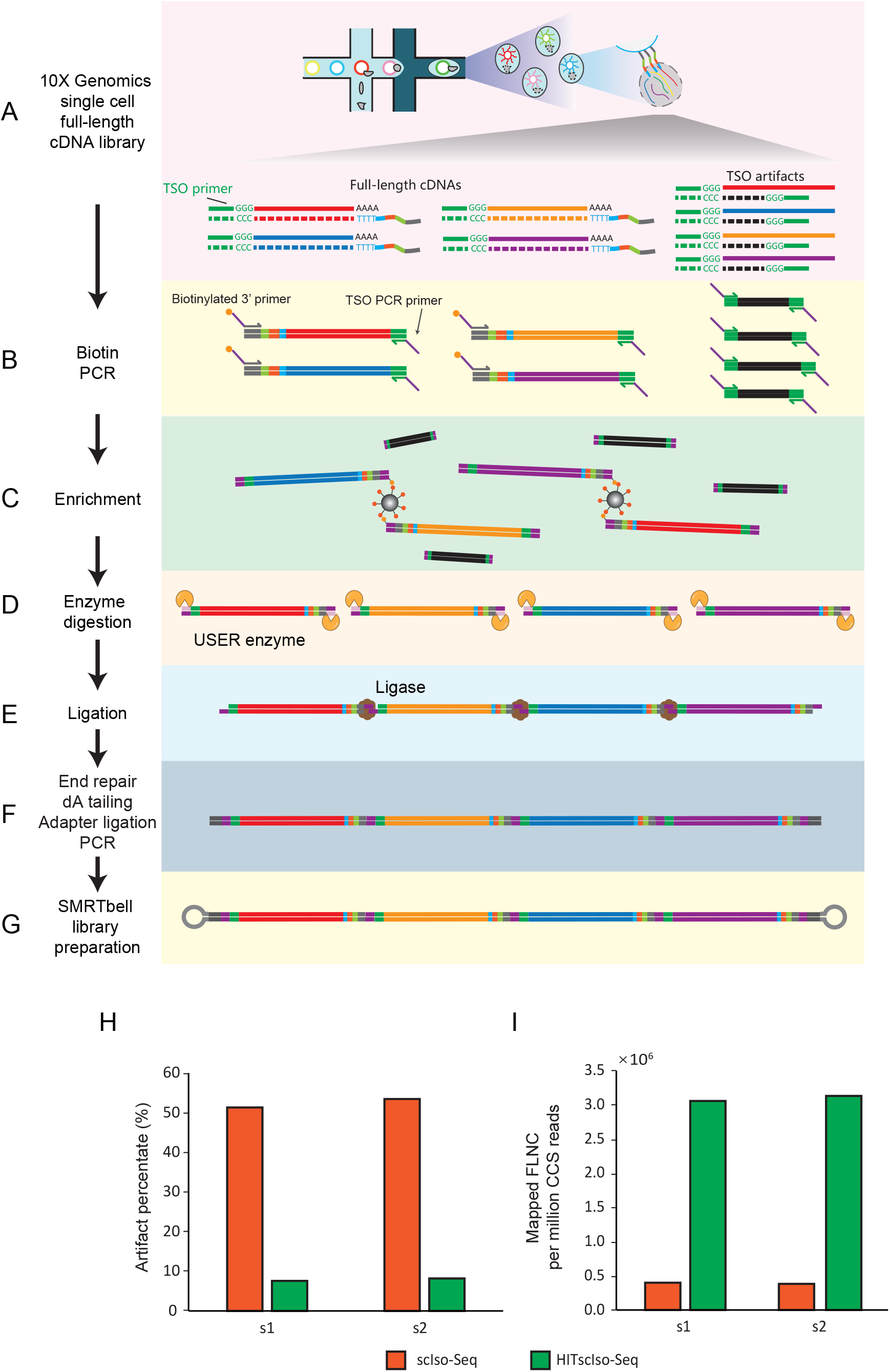
HIT-scISOseq workflow and performance. (A) Overview of the 10X Genomics single-cell full-length cDNA library construction. (B) PCR reaction is used to amplify full-length cDNAs with a biotinylated 3’ primer. (C) The biotinylated full-length cDNAs are enriched by streptomycin magnetic beads. (D) Restriction digestion at both ends to produce sticky end by USER enzyme. (E) Ligation of the cDNAs. (F-G) End repair, dA tailing, adapter ligation, and PCR enrichment of the ligated cDNAs, followed bySMRTbell library preparation. (H) Comparison of percentages of artifact reads between scISOseq and HIT-scISOseq. (I)Comparison of the amount of Mapped FLNC reads between scISOseq (orange) and HIT-scISOseq (green).

### Performance of HIT-scISOseq

To compare sequencing read output from different library preparation methods using the same PacBio Sequel II instrument and Sequel II SMRT cell, we evaluated the following methods: scISOseq (Figure S1), Linked-scISOseq (Figure S2), and HIT-ISOseq (Figure 1). Among these, scISOseq is a standard library preparation method (without involving capture and concatenation procedures) that is widely used and for which we had the expertise to prepare CCS2 libraries. Linked-scISOseq differs from HIT-scISOseq in that it includes only a full-length cDNA concatenation procedure, but no biotinylated primers are used to capture truly barcoded cDNAs. A comparison of the three methods allows us to assess each procedure’s relative performance. We directly compared the performance of the three methods using the same RNA samples from limbal epithelium, each sample with two replicates (s1 and s2), for which the transcriptional profiles have been well characterized previously. A total of five SMRTbell cells was sequenced on PacBio Sequel II system, as we only used s1 sample for Linked-scISOseq due to our budget prioritization. The libraries were sequenced following the ISO-Seq sample preparation protocol with recommended loading concentrations for submission (Supplementary Tables 1-4). As the computational analysis of concatenated full-length cDNAs requires special considerations due to the physical proximity of multiple cDNAs as well as the random 5’-to-3’ direction, we developed an isoform data analysis pipeline (scISA-Tools, see method sections) to enable identification and quantification of poly(A) tail, cell barcode (cellBC), and unique molecular identifier (UMI), and assignment of reads to the cell and RNA molecule. Based on PacBio’s recommended ISOseq data processing procedure, the mapped cDNAs were further classified as full-length non-chimeric (FLNC), non-full-length (NFL), and artifact reads, based on the presence of the poly(A) tail signal and the 5’ and 3’ cDNA primers. Artifact reads refer to the ones with neither 3’ primer nor poly(A) tail.

Performance assessment can be roughly divided into several process elements, including raw polymerase reads, CCS reads, FLNC reads and mapped FLNC reads (Table 1). At the level of raw polymerase reads, the yields of the three methods were similar (ranging from 4.30 M to 5.69 M), as with the percentage of productive ZMWs (P1 percentage metric, ranging from 53.75% to 71.13%). The similarity in polymerase reads suggests that the SMRTbell cDNA templates produced by the three methods were of high quality. Also, we found that all the methods had a long polymerase read length (> 70 kb), suggesting a good quality in the instrument run. Such a long polymerase length could theoretically allow for passing over a long insert of interest (typically peak at 4.9 kb to 5.2 kb in the current study) with a minimum three full passes and predicted accuracy of more than 0.9 (default requirement for CCS sequencing analysis). This is supported by our sequence data that the QV values could reach 0.97 with full passes ≥3. The average full passes were > 20 for both the linked-scISOseq and the HIT-scISOseq, indicating a high consensus accuracy for both methods. Notably, the polymerase read lengths generated from Link-scISOseq and HIT-scISOseq were 70% of that generated from standard scISOseq (Figure S4). It may be that the polymerases on more extended linked cDNA inserts have slightly lower survival, presumably due to DNA damage on linked inserts in SMRTbell libraries that hampers the polymerase reaction. Owing to the shorter polymerase read lengths, the yield of polymerase reads generated from Linked-scIOSseq and HIT-scISOseq was relatively lower than the number from scISOseq.

**Table1.** Performance of throughput and accuracy of scISOseq and HIT-scISOseq on two replicate samples of Cynomolgus monkey limbal cells.

|  | Sample | scISOseq |  | Linked-scISOseq | HIT-scISOseq |  |
| --- | --- | --- | --- | --- | --- | --- |
|  |  | s1 | s2 | s1 | s1 | s2 |
| Raw Data<sup>a</sup> | Polymerase reads count (M) | 4.95 | 4.30 | 5.02 | 4.74 | 5.69 |
|  | Yield of polymerase reads (GB) | 499.77 | 415.52 | 365.12 | 383.74 | 438.64 |
|  | Avg. polymerase reads length (Kb) | 101.06 | 96.66 | 72.78 | 80.88 | 77.06 |
|  | Yield of subreads (GB) | 487.53 | 405.99 | 361.45 | 379.87 | 434.44 |
|  | Avg. subreads length (Kb) | 1.55 | 1.68 | 3.64 | 3.46 | 3.61 |
| CCS Reads<sup>b</sup> | CCS reads count (M) | 4.02 | 3.38 | 3.70 | 3.43 | 4.23 |
|  | Yield of CCS reads (GB) | 8.04 | 7.12 | 16.56 | 16.75 | 21.62 |
|  | Avg. CCS reads length (Kb) | 2.00 | 2.11 | 4.48 | 4.89 | 5.11 |
|  | Avg. CCS reads passes | 70 | 64 | 21 | 23 | 20 |
|  | Avg. CCS reads QV | 0.97 | 0.97 | 0.95 | 0.95 | 0.95 |
| FLNC Detection<sup>c</sup> | Linked cDNA count (M) | 3.44 | 2.90 | 11.57 | 11.64 | 14.84 |
|  | FLNC count (M) | 1.60 | 1.29 | 5.25 | 10.47 | 13.23 |
|  | NFL count (M) | 0.07 | 0.06 | 0.14 | 0.28 | 0.39 |
|  | Artifact RNA count (M) | 1.76 | 1.55 | 6.18 | 0.88 | 1.22 |
|  | FLNC percentage (%) | 46.59 | 44.53 | 45.34 | 89.99 | 89.15 |
|  | NFL percentage (%) | 2.10 | 2.03 | 1.24 | 2.43 | 2.66 |
|  | Artifact RNA percentage (%) | 51.31 | 53.44 | 53.42 | 7.58 | 8.20 |
| FLNC Mapping<sup>d</sup> | Mapped FLNC Count (M) | 1.59 | 1.29 | 5.20 | 10.34 | 13.05 |
|  | Mapped FLNC percentage (%) | 99.44 | 99.50 | 99.05 | 98.75 | 98.65 |
|  | Avg. FLNC mapping coverage (%) | 99.13 | 99.14 | 98.88 | 98.90 | 98.83 |
|  | Avg. FLNC mapping identity (%) | 98.45 | 98.39 | 97.60 | 97.74 | 97.59 |
|  | Avg. collapsed FLNC length (Kb) | 2.37 | 2.47 | 2.18 | 2.22 | 2.24 |

**a**, the rows show (from up to down): (i) total polymerase reads count (million) of each sample; (ii) sum of all polymerase reads bases (gigabase) in each sample; (iii) average polymerase reads length (kilobase) in each sample; (iv) sum of all subreads bases (gigabase) in each sample; (v) average subreads length (kilobase) in each sample.
**b**, the rows show (from up to down): (i) total CCS reads count (million) of each sample; (ii) sum of all CCS reads bases (gigabase) in each sample; (iii) average CCS reads length (kilobase) in each sample; (iv) average CCS reads passes in each sample; (v) average CCS reads QV (Phred 33) in each sample.
**c**, the rows show (from up to down): (i) total linked cDNAs (which defined as linked cDNA in each CCS reads) count (million) in each sample; (ii) total full-length non-concatemer (FLNC) reads count (million) in each sample; (iii) total non-full length (NFL) reads count (million) in each sample; (iv) total artifact cDNA count (million) in each sample; (v) percentage of FLNC in linked cDNAs of each sample; (vi) percentage of NFL in linked cDNAs of each sample; (vii) the percentage of artifact cDNAs in linked cDNAs of each sample.
**d**, the rows show (from up to down): (i) total mapped FLNC count (million) of each sample; (ii) percentage of mapped FLNC in total FLNC of each sample; (iii) average mapping coverage of mapping FLNC in total FLNC of each sample; (iv) average
mapping identity of mapping FLNC in total FLNC of each sample; (v) average collapsed FLNC reads (which mean the reads after mapping quality filter and collapsing of redundancy) length (kilobase) in each sample.

At the level of CCS reads, the performance was quite similar among the three methods (ranging from 3.38 M to 4.23 M, Table 1), and the CCS reads were positively associated with polymerase read counts. As expected, Linked-scISOseq and HIT-scISOseq generated longer average CCS read lengths (4.48 kb, Linked-scISOseq; 4.89 kb for S1 sample and 5.11 kb for S2 sample, HIT-scISOseq), which are more than twice of that from scISOseq (Figure S5). The average CCS read lengths were positively correlated with the lengths of concatenated full-length transcript during library construction (Figure S3). Despite the differences in CCS read length, the three methods generally had similar CCS read QV values (> 0.95), suggesting that high-quality CCS reads could be generated from long insert SMRTbell templates by ligation of multiple cDNA fragments.

At the level of FLNC reads, linked cDNAs were obtained after demultiplexing the sequence data using our scISA-Tools (see Methods section). Both Linked-scISOseq and HIT-scISOseq had a higher amount of linked cDNAs when compared with scISOseq (Table 1). It is noteworthy that HIT-scISOseq had a much lower level of artifact cDNA reads (7.58%, S1 sample; 8.20%, S2 sample) when compared with Linked-scISOseq (53.42%) and scISOseq (51.31%, S1 sample; 53.44%, S2 sample). This result indicates that the capture procedure in HIT-scISOseq is effective in removing the majority of artifact reads resulting from TSO-flanked fragments during library construction, and ultimately increasing the read yields. By excluding the impact of artifact reads, the net percentage of FLNC reads (FLNC/(NFL+FLNC)) was quite similar among the three methods.

At the level of mapped FLNC reads, the performance of HIT-scISOseq rivals that of scISOseq and Linked-scISOseq. After alignment of the PacBio reads against the monkey reference genome, HIT-scISOseq had the highest mapped reads (10.34 M, S1 sample; 13.05 M, S2 sample), providing 6x and 10x more reads per SMRTBell cell compared to scISOseq, and up to 2x more reads compared to Linked-scISOseq (Table 1). As a rough order of magnitude, our capture and concatenation procedures increased the mapped FLNC reads by factors of 2 and 4, respectively, with a combined yield increasing 8-fold. Despite the difference in read yield, the three methods showed similar mappability of FLNC reads. More than 98% was mappable with HIT-scISOseq, and more than 99% was mappable with standard scISOseq. The mappability again confirms the high quality of FLNC reads. The average FLNC mapping coverage values were > 98%, and the mapping identity values were > 97%, for both Linked-scISOseq and HIT-scISOseq, comparable to those generated from scISOseq. These mapping metrics confirm the robustness of our scISA-Tools pipeline for precise read alignment.

After mapping reads and collapsing identical alignments, we found that the average lengths of collapsed reads (transcripts) from HIT-scISOseq were comparable to those from scISOseq (Table 1, Figure S6). Therefore, any potential bias of transcript length distribution was not observed in the data obtained from HIT-scISOseq. Our study also provides a detailed comparison of the two biological replicate datasets generated from HIT-scISOseq. Results between replicates were consistent, confirming that HIT-scISOseq provides an 8-fold higher read yield per run with high accuracy at an ever-decreasing price compared with scISOseq. The subtle difference in read yield metrics between biological replicates may be due to the differences in the percentage of productive ZMW loading and in sample quality.

### Multi-level validation of HIT-scISOseq

One of the prominent uses of single-cell RNA sequencing is the quantification of gene expression and identification of distinct cell types by clustering scRNA-seq data. Given that the limbal epithelium consists of several well-defined cell types, we used limbal cDNA samples for cross-platform validation of HIT-scISOseq’s ability to distinguish different cell types. We compared HIT-scISOseq and Illumina short-read RNA sequencing (NGS) using the same single-cell 10xGenomics cDNA samples, and we identified a strong concordance between the two platforms. To begin with, we used HIT-scISOseq data to quantify gene expression using our scISA-tools pipeline. We found that there was a strong correlation between HIT-scISOseq and NGS in terms of UMI counts by cellBC (Pearson’s r > 0.990, p < 0.001, Figure 2A and Figure S7) and UMI counts by gene (Pearson’s r > 0.950, p < 0.001, Figure 2B and Figure S8). For the HIT-scISOseq datasets, there was a high agreement in UMI counts by gene in the two biological replicates (Figure 2C). Also, UMAP projection of gene expression data from the two platforms showed consistent results in cell-type classification (4 cell clusters, Figure 2D-F, Figure S9 and Supplementary Table 5) with clear cell type boundaries, including conjunctival cells, limbal cells, central basal cells, and differentiated cells. Comparing NGS and HIT-scISOseq gene expression quantification for the same cell type showed a high correlation (Pearson’s r > 0.95, Figure 2G and Figure S10), as with the percentage of shared cell barcodes for the same cell type (82.18% to 97.63%, Figure 2F). The high percentage of sharing in cellBC suggests that the HIT-scISOseq represents well the transcriptome identified in the 10x Genomics system. We next created heatmaps of the top 15 genes per cell cluster (Figure 2H-I and Figure S11). Data from the two platforms showed similar patterns of marker gene expression. These results confirm that the gene expression data derived from HIT-scISOseq recapitulate NGS gene expression quantification.

**Figure 2.**
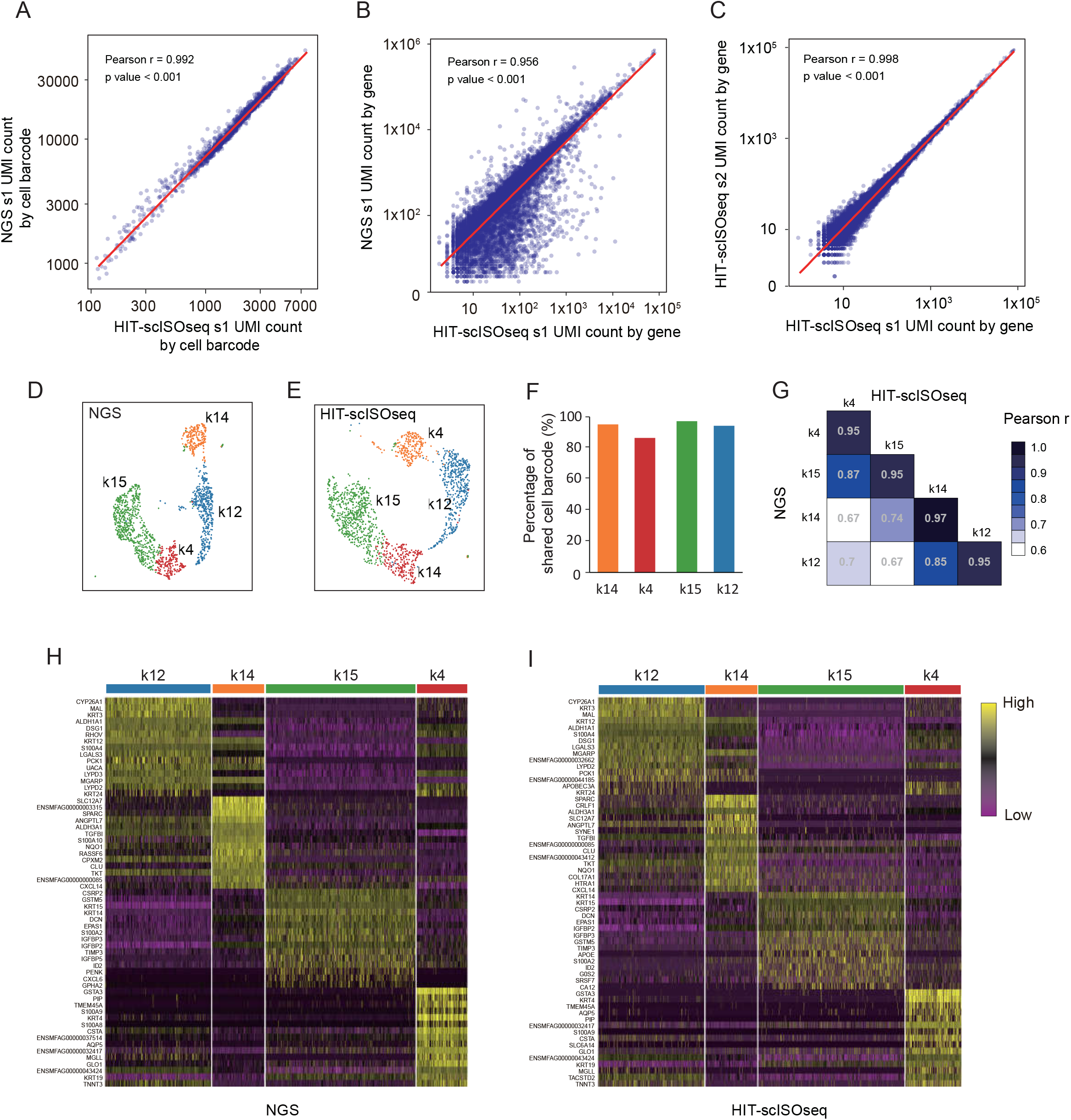
Accuracy and reproducibility of HIT-scISOseq. (A) Both NGS and HIT-scISOseq datasets are generated using the same limbal cDNA samples. Correlation dot plot of NGS (y-axis) and HIT-scISOseq (x-axis) UMI counts by cell barcode is shown. Correlation coefficient Pearson r, and p-value are shown. (B) Correlation dot plot of NGS (y-axis) and HIT-scISOseq (x-axis) UMI count by gene is shown. (C) Correlation dot plot of two HIT-scISOseq biological replicate samples UMI count by gene is shown. For each correlation dot plot, the x- and y-axis for count numbersareon a log10 scale. Gene expression profiles are determined independently for each cell cluster using either NGS or HIT-scISOseq. Both NGS (D) and HIT-scISOseq (E) data sets show that the four main cell clusters could be successfully clustered (differentiated cells (K12+) = blue, corneal basal cells (K14+) = orange, limbal stem cells (K15+) = green, and conjunctival cells (K4+) = red). (F) Bar charts showing the percentage of cell barcodes shared in the NGS and HIT-scISOseq data sets. (G) Correlation heatmap for gene expression in each cell cluster between NGS and HIT-scISOseq data sets. (H and I) Marker gene expression heatmaps of the four major cell clusters in NGS and HIT-scISOseq data sets. Color gradient represents log-transformed, and normalized counts scaled to a maximum of 1 per row. Upper bars represent cell clusters assignment for individual cells.

### Accurate isoform quantification by HIT-scISOseq

We next sought to identify and quantify single-cell isoforms using HIT-scISOseq datasets. Based on isoform-level information, we were able to create similar clustering of cells at the same pattern as gene-level clustering analysis (Figure 3A and Figure S12). Also, isoform-level expression was highly correlated between the two biological replicate samples. Further analysis of the top 15 marker isoforms of each cell type showed that up to half of these isoforms are previously annotated and known, whereas the rest of isoforms could be classified as novel isoforms, including known gene’s novel isoforms and novel gene’s isoforms (Figure 3B-C and Figure S13-14). Among the cell types identified, differentiated cells were the most differentiated subpopulation, which had a higher number of isoforms than the other cell types (Figure 3D and Figure S15), suggesting an active transcriptional process during cell differentiation. HIT-scISOseq data also enables the analysis of multiple isoforms of gene locus. For example, volcano plots across limbal and basal cell types allowed us to visualize the results of differential expressed isoforms (Figure 3F). Krt15 has been well documented as a marker gene for limbal cells, and isoforms derived from Krt15 varied in their expression level among different cell types (Figure 3G). These results suggest that HIT-scISOseq provides an ability to delineate isoform information with a much higher resolution that has been previously unavailable with short-read sequencing.

**Figure 3.**
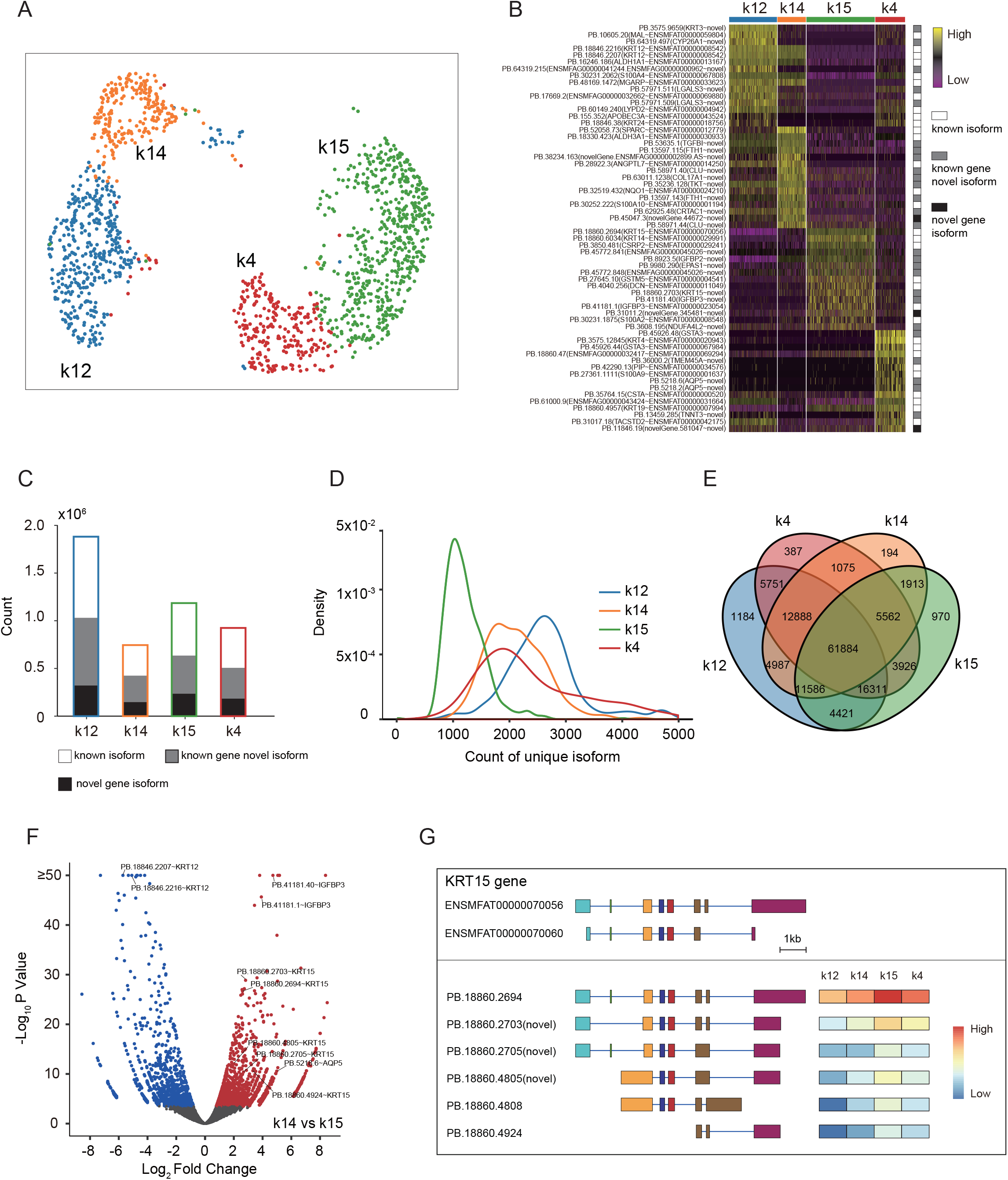
Single-cell isoform-level analysis highlights the heterogeneity of the limbal cell population. (A) Cell type clustering derived from HIT-scISOseq’s full-length isoform expression matrix data. The cellscould be successfully clustered (differentiated cells (K12+) = blue, corneal basal cells (K14+) = orange, limbal stem cells (K15+) = green, and conjunctival cells (K4+) = red). (B) Single-cell maker isoform expression heatmap showing cell-type-specific isoform marker expression among different cell clusters. Color gradient represents log-transformed, and normalized counts scaled to a maximum of 1 per row. Upper bars represent cell groups’ assignment for individual cells. Right panel color-coding bar shows known isoforms (white), known gene’s novel isoforms (grey), and novel gene’s isoforms (black). (C) Distribution of known and novel isoforms in each cell cluster. (D) Distribution of expressed unique isoform numbers (isoforms with minimal UMI count of 3) of individual cells in each cell cluster. (E) Venn plot of cell-type-specific isoforms.Y-axis for density of cell barcode count. (F) Volcano plot showing differential expression of isoforms between the corneal basal cell and limbal stem cell populations (Red and blue dot, FDR < 0.01 and |log2 Fold Change| > 0). (G) Differential usage of known and novel isoforms of KRT15 in different cell clusters. Top panel shows the reference annotation of known KRT15 isoforms. Bottom left panel shows the top 6 differentially expressed isoforms identified from the analysis in Figure 3F. Bottom right panel is an expression heatmap of these isoforms in the four cell clusters.

## Discussion

We show that HIT-scISOseq is a high-throughput and high accuracy method that could be used to characterize isoforms in thousands of nonhuman primate cells with very high accuracy. We demonstrate the ability of a scISA-tool computation pipeline to decode transcripts that could not be identified by short-read scRNA-seq alone. Despite this progress, there remains room for improving the performance of HIT-scISOseq. The quality and accuracy of long-read sequencing continue to evolve. The recent release of PacBio Sequel II SMRT Cell 8 M has allowed for long insert reads (15 kb) with high consensus accuracy (>99.9% for HiFi reads). Further advancements in sequencing chemistries and ZMW design will vastly improve the sequencing capacity for even longer reads. However, none of the current isoform sequencing technology has kept up with the pace of the PacBio’s Sequel II long-read sequencing capacity. Based on HIT-scISOseq, we discovered that, as the length of multiple concatenated cDNAs grew, there was an increased risk of irreversible cDNA damage (e.g., nicks) that could severely impair the performance of DNA polymerase in ZWMs. We have thus optimized our system by identifying 5 kb (though concatenation of 2 to 3 full-length transcripts) as the appropriate length for preparing long-insert libraries since this moderate level of concatenation does not lead to a high percentage of nicks. Further strategies to optimize the input quality by reducing DNA nicks would enable the construction of high-quality long inserts with more cDNA concatemers, and ultimately improving PacBio’s sequencing throughput. A modified workflow that includes size selection via BluePippin system to enrich for longer concatenated molecules might also be worthwhile for increasing the long-read yield.

While we have focused on mRNA transcripts, any transcripts of interest (e.g., LncRNA, Poly(A) tail, or eccRNA) can be targeted and enriched by altering the composition of concatenation libraries^17, 18^. Also, the high quality of full-length transcript will be applicable for the identification of both transcriptional information and somatic mutations at the single-cell level. The increased accuracy will allow for more straightforward phasing of transcripts at the single-cell level and allele-specific expression (ASE) analysis^19^. We also envision the use of HIT-scISOseq in multiplexed single-cell RNA-sequencing of pooled unrelated individuals, in which natural polymorphisms in long transcripts can be utilized to demultiplex reads and recover sample identify^6^. In addition, the increased accuracy allows for more direct alignment, reducing the need for error correction using NGS information. Finally, we have shown that the HIT-scISOseq is fully compatible with a commercially available single-cell platform (10x Genomics), and it should be readily adaptable to other microwell-based and combinatorial-indexing-based technologies. In conclusion, the availability of low cost, high throughput HIT-scISOseq represents an advance in single-cell technology that will allow the power of population-scale analysis to be realized.

## Methods

### Monkey limbal wounding experiment

All animal experiments were conducted in accordance with the ARVO Statement for the Use of Animals in Ophthalmic and Vision Research and approved by the institutional animal care and use committee of Zhongshan Ophthalmic Center, Sun Yat-sen University. Cynomolgus monkeys (Macacafascicularis) were anesthetized with a mixture of ketamine and xylazine, and topical anesthesia with 0.5% proparacaine hydrochloride (Alcaine; Alcon). Only female monkeys aged four years were used. Whole circumference (360-degree) limbal excision was performed in theeye. The limbal excision was made by lamellar dissection of the limbal zone, 2 mm into the cornea and 2 mm into the conjunctiva, and 100 um in depth. Wounds were left to heal by secondary intention, and punch biopsies were made at 0 (healthy), 3, and 48 hours for single-cell experiments. Biopsy tissues were placed into cryovials with Advanced DMEM F-12 and placed on ice.

### Single-cell dissociation

Dissected limbal tissue was micro-dissected and disaggregated into single cells with Dispase II (Sigma) and Collagenase IV (Sigma) at 37°C with constant rotation. The epithelial layer was isolated from the underlying stroma and separately digested at 37°C for 2 hours with 2 mL of 1 mg/mL collagenase A (Sigma-Aldrich Corp., St. Louis, MO, USA) in Dulbecco’s modified Eagle’s medium (DMEM) containing 10% FBS, 50 μg/mL gentamicin, and 1.25 μg/mL amphotericin B. Clusters were further digested with 0.25% trypsin and one mM EDTA with gentle pipetting to yield single cells. Approximately 10,000 epithelial cells and 10,000 stromal cells were pooled in order to recover sufficient numbers of epithelial and stromal cells for downstream analyses. The mix cells were filtered through a 30-um cell strainer, and re-suspended in 60 uL PBS containing 0.04% BSA to achieve 1,000 cells/ul for capture on the 10x Genomics Chromium controller.

### 10x Genomics single-cell capture and Illumina library preparation

The dissociated single cells were processed through the GemCode Single Cell Platform per manufacturer’s recommendations using the Chromium Single Cell 3’ GEM, Library, and Gel Bead Kit v3 (10x Genomics; PN-1000075) with a recovered amount of approximately 2,000 cells. Illumina library preparation was performed using Chromium Single Cell 3’ Reagent Kits User Guide (V3 Chemistry). After the cDNA cleanup step (Step 2.1), half of the purified cDNA was used for PacBio library preparation, whereas the rest was used for downstream Illumina library preparation. Illumina libraries were sequenced on a NextSeq 550 (SY-415-1002, Illumina) using the NextSeq High Output Kit (150 cycles; 20024907, Illumina) with the following read protocol: read 1, 118 cycles; i7 index read, 8 cycles; read 2, 40 cycles.

### cDNA amplification and capture for PacBio library construction

cDNA products were amplified with KAPA HiFi HotStart Uracil 2 x ReadyMix (Kapa Biosystems) and newly designed PCR primers that contain a deoxyuraciland one of which was biotinylated. PCR products were then purified with AgencourtAMPure XP Beads (Beckman Coulter), quantified with Qubit dsDNA HS Assay Kit (Thermo Fisher), and assessed with Agilent 2100 DNA HS Assay (Figure S3). The barcode-UMI-Poly(dT)-flanked cDNAswerecaptured on the streptavidin-coated M-280 dynabeads using Dynabeads™ kilobaseBINDER™ Kit (60101, Invitrogen, Carlsbad, CA), whereas the unbound cDNAs were removed.

### USER cloning-based ligation of multiple inserts

cDNA products on the dynabeads were washed with wash buffer and nuclease-free water, re-suspended with 19 μl reaction buffer contain 2ul 10x T4 DNA ligase buffer (NEB), 1 μl USER Enzyme (NEB) and incubated at 37 °C for 20 min, nicking the deoxyuracil site to generates 3’ palindrome overhangs suitable for ligation of multiple inserts and simultaneously release the cDNA from M-280 dynabeads. One microlitre of T4 DNA ligase (NEB, 400,000U/mL) was added to the reaction and incubated at 16 °C to ligate inserts. The resultant multi-insert library was purified withAgencourtAMPure XP Beads (Beckman Coulter) and then end-repaired and A-tailed using the NEBNext Ultra II End Repair/dA-Tailing Module for 15 min at 20 °C and then 30 min at 65 °C. The cDNA was ligated with a dT-overhang selection adapter using the NEBNext^®^ Ultra™ II Ligation Module (NEB) for 15 min at 20 °C, purified with AgencourtAMPure XP Beads (Beckman Coulter), and PCR amplified using KAPA HiFi HotStart 2x ReadyMix to screen out the multi-insert library without ligation nicks. The amplified products were again purified by AgencourtAMPure XP Beads (Beckman Coulter) and assayed with Agilent DNA 12000 Assay.

### PacBio SMRTbell templates preparationandsequencing

Amplified PCR products were end-repaired and A-tailed using NEBNext End Repair/dA-Tailing Module, ligated with a dT-overhang hairpin adapter using the NEBNext^®^ Ultra™ II Ligation Module (NEB), and purified with AgencourtAMPure XP Beads (Beckman Coulter) to produce the SMRTbell Template. To remove residual adapters and unligated DNA fragments, we added 1 μlExonuclease I (NEB), 1 μl Exonuclease III (NEB), and NEBuffer 1 (NEB) to the library and incubated at 37 °C for 1 hour. The products were purified using AgencourtAMPure XP Beads and quantified using the Agilent DNA 12000 Kit (Agilent). Sequencing primer annealing and polymerase binding to the PacBio SMRTbell Templates were performed using the manufacturer’s recommendations (PacBio, US).Library complex wasthen sequenced using SMRT Cell 8M (PacBio)compatible with the Sequel II sequencer.

### HIT-scISOseq data processing pipeline

Since HIT-scISOseq links multiple transcripts together, and multiple cDNA-library-prep-primer sequences could be found in one CCS read, the PacBio official IsoSeq3 pipeline would inherently define HIT-scISOseq reads as “chimeric” and thus the pipeline is not suitable for our analysis. Here, we have developed a set of analysis tools (https://github.com/shizhuoxing/scISA-Tools) as a pipeline for 10X Genomics scISOseq reads processing. This pipeline includes quality control, basic statistics, full-length transcripts identification, cell barcode and UMI extraction and correction, isoform clustering, single-cell isoform quantification, and single-cell expression matrix format transform (Supplementary Table 2). This pipeline is not only useful for HIT-scISOseq data. It also works well in 10X Genomics based on standard scISOseq protocol.

### API for interactive visualization of Loupe Browser

Loupe Browser is an established desktop application that provides interactive visualization of single-cell RNA data from 10X Genomics platform. We have developed scMatrix2CellRangerH5, a utility that could convert text matrix to h5 format that is compatible with the CellRanger reanalyze pipeline so that cloupe files could be generated and visualized in Loupe Browser.

### Single-cell short read analysis

For each sample, we used the 10X Genomics CellRanger pipeline (version 3.1.0) to obtain a single-cell expression matrix base on Macacafascicularis genome and transcriptome (Ensembl Macaca_fascicularis_5.0.99).

### Single-cell isoform sequencing and bioinformatics pipeline

#### Generation of Circular Consensus Sequencing Reads

Using SMRT-Link (version 8.0.0.80529), we generated CCS reads with the following modified parameters: “--min-passes 0 --min-length 50 --max-length 21000 --min-rq 0.75”.

#### Generation of Single Cell Full-Length Non-Concatemer (FLNC) Reads

First, we mapped the 5’ and 3’ primers to CCS reads using NCBI BLAST (version 2.10.0+)^20, 21^with parameters: “-outfmt 7 -word_size 5”. We then considered primer blast results as inputs, using the classify_by_primer utility to extract cell barcodes and UMIs, and then generated FLNC with the following parameters: “-min_primerlen 16 -min_seqlen 50”. The details of classify_by_primer utility are briefly listed as follows: (1) parsing standard pair of 5’ and 3’ primers in CCS reads to obtain full-length isoforms, which were then oriented from 5’ to 3’ end; (2) trimming 5’ and 3’ primer sequences, followed by trimming the 28bp sequences in the 3’ primers as cell barcodes and UMIs; and (3) trimming 3’ polyA tail using a sliding window algorithm. As our program was strictly 5’ and 3’ primer paired one after another, each full-length read was oriented. The reads with primers, cellBCs, UMIs, and polyA tails, were considered as full-length non-concatemer (FLNC) reads.

#### Genome alignment of FLNC reads

After FLNC detection and trimming procedures to obtain primers, cellBCs, UMIs and polyA tails, the remaining fraction of each FLNC was aligned to the Macacafascicularis genome (Ensembl Macaca_fascicularis_5.0.99) with minimap2 (version 2.17-r974-dirty)^22^ in spliced alignment mode with the following parameters: “-ax splice -uf --secondary=no -C5”.

#### Cell Barcode and UMI correction

We adopted a strategy similar to the one for 10X Genomics CellRanger. We warped cellBC correction function in CellRanger as a module in our pipeline named cellBC_UMI_corrector. This utility could handle long read data without the need to relate to short-read information for guiding.

For cellBC correction, CellRanger based on known barcodes for given assay chemistry were stored in an “allowlist” file. The steps are briefly described as follows:

1. We counted the observed frequency in the dataset of every barcode on the allowlist
2. For every observed barcode that is 1-Hamming distance (substitution) away from theallowlist, we computed the posterior probability that the observed barcode originated from the allowlist barcode with a sequencing error at the differing base (based on the base Q score). We next replaced the observed barcode with the allowlist barcode with the highest posterior probability that exceeds 0.975

For UMI correction, the steps are briefly described as follows:

1. Performs some basic quality filtering and correction for UMI sequencing errors

a. Must not be a homopolymer, e.g. AAAAAAAAAA;
b. Must not contain N;
c. Must not contain bases with base quality < 10.
2. UMIs that are 1 Hamming distance (substitution) away from a higher-count UMI were corrected to the higher count UMI if they shared a cell barcode

#### Generation of single-cell gene count matrix

After mapping FLNCs to the genome, we used a gffcompare (version 0.11.6)^23^ and assigned the FLNCs to Ensemble Macacafascicularis annotation gene models (Ensembl Macaca_fascicularis_5.0.99). The reads were defined as exonic sequences when the class codes equaled to the “= c k m n j e o”. This procedure is consistent with the CellRanger pipeline. Next, we used scGene_matrix utility to generate the single-cell gene expression data for each sample, based on gffcompare output and corrected cellBC and UMI on each FLNC.

#### Collapsing redundant isoforms

We used cDNA_Cupcake (https://github.com/Magdoll/cDNA_Cupcake) python script collapse_isoforms_by_sam.py. The default settings --min-coverage for minimum alignment coverage and --min-identity for minimum alignment identity were 0.99 and 0.95, respectively. This step is to ensure the generation of transcripts with high accuracy.

#### Non-redundant isoforms classification, coding frame prediction, and UTR detection

We used SQANTI3 (https://github.com/ConesaLab/SQANTI3)^24^ to characterize non-redundant isoforms based on Ensemble Macacafascicularis annotation gene models (Ensembl Macaca_fascicularis_5.0.99). Isoforms were classified as known or novel ones. SQANTI3 was used to call GeneMarkS-T (version 5.1 March 2014) for non-redundant isoforms CDS coding frame prediction and UTR regions definition.

#### Generation of single-cell isoform count matrix

After the collapsing procedure, we used scIsoform_matrixutility to generate single-cell isoform expression quantity in each sample with the following parameters: “-minUMIcount 3”.

#### Expression matrix’s quality control

We used the Seurat R package (version 3.1.5)^25^ and performed quality filtering analysis of each sample’s single cell gene and isoform expression matrix. We used “min.cells = 5, nFeature_RNA> 200, nFeature_RNA< 6000, percent.mt < 25” for NGS B0h1 and B0h2 gene expression matrix, “min.cells = 5, nFeature_RNA> 100, nFeature_RNA< 3000, percent.mt < 25” for TGS B0h1 gene expression matrix, “min.cells = 5, nFeature_RNA> 100, nFeature_RNA< 3500, percent.mt < 25” for TGS B0h2 gene expression matrix, and “min.cells = 5, nFeature_RNA> 100, nFeature_RNA< 4500, percent.mt < 25” for TGS B0h1 and B0h2 isoform expression matrix.

#### Cell clustering and cell-type annotation

After the quality filtering procedure, we used scMatrix2CellRangerH5 utility to convert the matrix to CellRanger h5 format, and then we used CellRanger reanalyze pipeline for PCA and cell clustering with default parameters. The resulting cloupe files were loaded to Loupe Browser for fine manual annotation of cell types and tune adjustments. After cell-type annotation, we imported the cell type and cell barcode associated tables to the Seurat R package (version 3.1.5) for downstream cell clustering, as well as cell-type marker-gene and marker-isoform expression heatmap generation.

#### Differential expression analysis of Gene and Isoform

We used tappAS Application (version 1.0.1)^26^ for cell-type gene and isoform differential expression analysis.

#### Generation of Isoforms structure view

**Selected interested isoforms were imported as GTF files to IGV (version 2.8.2)^27^ for splicing structure viewing.**

## Reference

1. Tang, F.C. et al. mRNA-Seq whole-transcriptome analysis of a single cell. Nat Methods6, 377–U386 (2009).

2. Saliba, A.E., Westermann, A.J., Gorski, S.A. & Vogel, J. Single-cell RNA-seq: advances and future challenges. Nucleic Acids Research42, 8845–8860 (2014).

3. Fuccillo, M.V. et al. Single-Cell mRNA Profiling Reveals Cell-Type-Specific Expression of Neurexin Isoforms. Neuron87, 326–340 (2015).

4. Petropoulos, S. et al. Single-Cell RNA-Seq Reveals Lineage and X Chromosome Dynamics in Human Preimplantation Embryos. Cell165, 1012–1026 (2016).

5. Seow, J.J.W., Wong, R.M.M., Pai, R. & Sharma, A. Single-Cell RNA Sequencing for Precision Oncology: Current State-of-Art. Journal of the Indian Institute of Science, 1 (2020).

6. Kang, H.M. et al. Multiplexed droplet single-cell RNA-sequencing using natural genetic variation. Nat Biotechnol36, 89 (2018).

7. Arzalluz-Luque, A. & Conesa, A. Single-cell RNAseq for the study of isoforms-how is that possible? Genome Biol19 (2018).

8. Hardwick, S.A., Joglekar, A., Flicek, P., Frankish, A. & Tilgne, H.U. Getting the Entire Message: Progress in Isoform Sequencing. Front Genet10 (2019).

9. Picelli, S. et al. Full-length RNA-seq from single cells using Smart-seq2. Nat Protoc9, 171–181 (2014).

10. Lebrigand, K., Magnone, V., Barbry, P. & Waldmann, R. High throughput, error corrected Nanopore single cell transcriptome sequencing. BioRxiv, 831495 (2019).

11. Hagemann-Jensen, M. et al. Single-cell RNA counting at allele and isoform resolution using Smart-seq3. Nat Biotechnol38, 708–+ (2020).

12. Volden, R. & Vollmers, C. Highly Multiplexed Single-Cell Full-Length cDNA Sequencing of human immune cells with 10X Genomics and R2C2. BioRxiv (2020).

13. Wang, X. et al. Full-length transcriptome reconstruction reveals a large diversity of RNA and protein isoforms in rat hippocampus. Nature Communications10, 1–15 (2019).

14. Wenger, A.M. et al. Accurate circular consensus long-read sequencing improves variant detection and assembly of a human genome. Nat Biotechnology37, 1155–1162 (2019).

15. Elizabeth Tseng et al. Single cell isoform sequencing (scIso-Seq) identifies novel full-length mRNAs and cell type-specific expression. https://www.pacb.com/wp-content/uploads/Tseng-ENCODE-2019-Single-Cell-Isoform-Sequencing-scIso-Seq-Identifies-Novel-Full-length-mRNAs-and-Cell-Type-Specific-Expression.pdf. (2019).

16. Gupta, I. et al. Single-cell isoform RNA sequencing characterizes isoforms in thousands of cerebellar cells. Nat Biotechnol36, 1197–+ (2018).

17. Legnini, I., Alles, J., Karaiskos, N., Ayoub, S. & Rajewsky, N. FLAM-seq: full-length mRNA sequencing reveals principles of poly (A) tail length control. Nat Methods16, 879–886 (2019).

18. Liu, Y., Nie, H., Liu, H. & Lu, F. Poly (A) inclusive RNA isoform sequencing (PAIso− seq) reveals wide-spread non-adenosine residues within RNA poly (A) tails. Nature Communications10, 1–13 (2019).

19. Deonovic, B., Wang, Y., Weirather, J., Wang, X.-J. & Au, K.F. IDP-ASE: haplotyping and quantifying allele-specific expression at the gene and gene isoform level by hybrid sequencing. Nucleic Acids Research45, e32–e32 (2017).

20. Altschul, S.F., Gish, W., Miller, W., Myers, E.W. & Lipman, D.J. Basic local alignment search tool. Journal of Molecular Biology215, 403–410 (1990).

21. Camacho, C. et al. BLAST+: architecture and applications. BMC Bioinformatics10, 421 (2009).

22. Li, H. Minimap2: pairwise alignment for nucleotide sequences. Bioinformatics34, 3094–3100 (2018).

23. Pertea, G. & Pertea, M. GFF Utilities: GffRead and GffCompare. F1000Research9 (2020).

24. Tardaguila, M. et al. SQANTI: extensive characterization of long-read transcript sequences for quality control in full-length transcriptome identification and quantification. Genome Res28, 396–411 (2018).

25. Butler, A., Hoffman, P., Smibert, P., Papalexi, E. & Satija, R. Integrating single-cell transcriptomic data across different conditions, technologies, and species. Nat Biotechnol36, 411–420 (2018).

26. de la Fuente, L. et al. tappAS: a comprehensive computational framework for the analysis of the functional impact of differential splicing. Genome Biol21, 1–32 (2020).

27. Thorvaldsdóttir, H., Robinson, J.T. & Mesirov, J.P. Integrative Genomics Viewer (IGV): high-performance genomics data visualization and exploration. Briefings in Bioinformatics14, 178–192 (2013).

