## Supplementary Tables for "HIT-scISOseq: High-throughput and High-accuracy Single-cell Full-length Isoform Sequencing for Corneal Epithelium"

**Supplementary Table 1. Quality control reports of scISOseq, Linked-scISOseq, and HIT-scISOseq data sets**

| | scISOseq | | Linked-scISOseq | HIT-scISOseq | |
|---|---|---|---|---|---|
| Sample | s1 | s2 | s1 | s1 | s2 |
| Sequencing ID | 20PW00010 | 20PW00011 | 20PW00012 | 20PW00014 | 20PW00015 |
| Total Bases (GB) | 499.77 | 415.52 | 365.12 | 383.74 | 438.64 |
| Polymerase reads | 4,945,233 | 4,298,982 | 5,016,861 | 4,744,729 | 5,691,910 |
| Polymerase read N50 | 192,520 | 189,884 | 152,456 | 168,936 | 159,112 |
| Polymerase mean read length | 101,060 | 96,655 | 72,778 | 80,878 | 77,063 |
| Subreads bases (Gb) | 487.53 | 405.99 | 361.45 | 379.87 | 434.44 |
| Subreads number | 314,884,226 | 242,322,264 | 99,394,233 | 109,779,588 | 120,449,533 |
| Subreads mean read length | 1,548 | 1,675 | 3,637 | 3,460 | 3,607 |
| Ab 10k (%) | 5.74 | 6.57 | 12.95 | 12.62 | 12.55 |
| Unique molecular yield (GB) | 27.83 | 27.04 | 44.07 | 45.42 | 51.26 |
| P0 (%) | 35.73 (2,860,439) | 44.23 (3,540,019) | 33.72 (2,700,309) | 38.75 (3,104,726) | 25.54 (2,046,498) |
| P1 (%) | 61.88 (4,953,881) | 53.86 (4,310,507) | 62.71 (5,022,629) | 59.24 (4,746,738) | 71.06 (5,693,769) |
| P2 (%) | 2.50 (200,351) | 2.05 (164,145) | 3.64 (291,733) | 2.04 (163,207) | 3.42 (274,404) |
| CPolyReads number | 8,607 | 11,485 | 5,738 | 1,989 | 1,825 |
| CPolyReads mean length | 71,429 | 53,472 | 49,963 | 41,407 | 49,942 |
| CPolyConcordance | 0.85 | 0.84 | 0.84 | 0.84 | 0.83 |
| Data loss ratio (%) | 2.45 | 2.29 | 1.00 | 1.01 | 0.96 |

Total Bases (GB): sum of all polymerase reads bases (giga base) in each sample;
Polymerase reads: total polymerase reads count of each sample;
Polymerase N50 read length: 50% of all polymerase reads are longer than this value;
Polymerase mean read length: the mean read length of all polymerase reads;
Subreads bases (Gb): sum of all subreads bases (giga base) in each sample;
Subreads number: total subread count of each sample;
Subreads mean read length: the mean read length of all subreads;
Ab 10k (%): the percentage of subreads longer than 10kb;
Unique molecular yield (GB): sum of all unique molecular reads bases (giga base) in each sample;
P0 (%): The percentage of ZMWs that are empty, with no polymerase;
P1 (%): The percentage of ZMWs that are productive and sequencing;
P2 (%): The percentage of ZMWs that are not P0 (empty) or P1 (productive);
CPolyReads number: The number of control polymerase reads;
CPolyReads mean length: The mean polymerase read length of control reads;
CPolyConcordance: The average concordance (agreement) between the control raw reads and the control reference sequence;
Data loss ratio (%): The percentage of loss bases in generating reads from polymerase reads to subreads, i.e., the percentage of base of N or those low-quality bases in total bases.

**Supplementary Table 2. Cell barcode and UMI correction reports of scISOseq, Linked-scISOseq and HIT-scISOseq data sets**

| | | | scISOseq | | Linked-scISOseq | HIT-scISOseq | |
|---|---|---|---|---|---|---|---|
| | | Sample | s1 | s2 | s1 | s2 | s1 |
| | | FLNC read count | 1,602,410 | 1,291,533 | 5,245,279 | 10,472,456 | 13,227,218 |
| Cell barcode (CB) | All | CB in allowlist | 1,466,251 | 1,177,044 | 4,452,458 | 8,823,984 | 11,051,565 |
| | | CB in allowlist (%) | 91.50 | 91.14 | 84.89 | 84.26 | 83.55 |
| | | Corrected CB | 13,431 | 10,814 | 124,486 | 248,384 | 324,201 |
| | | Corrected CB (%) | 0.84 | 0.84 | 2.37 | 2.37 | 2.45 |
| | | Total number of corrected CB | 1,479,682 | 1,187,858 | 4,576,944 | 9,072,368 | 11,375,766 |
| | | Total number of corrected CB (%) | 92.34 | 91.97 | 87.26 | 86.63 | 86.00 |
| | CB QV>=0.95 (passed) | FLNC read count | 1,533,988 | 1,232,653 | 4,655,184 | 9,375,747 | 11,729,087 |
| | | CB in allowlist | 1,457,633 | 1,169,679 | 4,361,574 | 8,648,780 | 10,820,554 |
| | | CB in allowlist (%) | 90.97 | 90.57 | 83.15 | 82.59 | 81.81 |
| | | CB correction | 8,709 | 6,712 | 63,986 | 125,123 | 159,534 |
| | | CB correction (%) | 0.54 | 0.52 | 1.22 | 1.19 | 1.21 |
| | | Total number of corrected CB | 1,466,342 | 1,176,391 | 4,425,560 | 8,773,903 | 10,980,088 |
| | | Total number of corrected CB (%) | 95.59 | 95.44 | 95.07 | 93.58 | 93.61 |
| | CB QV>=0.99 | FLNC read count | 1,491,708 | 1,195,632 | 4,293,626 | 8,681,819 | 10,800,950 |
| | | CB in allowlist | 1,434,549 | 1,149,143 | 4,154,878 | 8,256,408 | 10,294,835 |
| | | CB in allowlist (%) | 89.52 | 88.98 | 79.21 | 78.84 | 77.83 |
| | | CB correction | 5,777 | 4,274 | 28,831 | 55,884 | 66,022 |
| | | CB correction (%) | 0.36 | 0.33 | 0.55 | 0.53 | 0.50 |
| | | Total number of corrected CB | 1,440,326 | 1,153,417 | 4,183,709 | 8,312,292 | 10,360,857 |
| | | Total number of corrected CB (%) | 96.56 | 96.47 | 97.44 | 95.74 | 95.93 |
| UMI | CB passed and UMI passed | Discarded UMI | 636 | 397 | 7,897 | 19,011 | 30,719 |
| | | Discarded UMI (%) | 0.04 | 0.03 | 0.15 | 0.18 | 0.23 |
| | | Uncorrected UMI | 1,465,304 | 1,175,682 | 4,414,536 | 8,743,058 | 10,931,494 |
| | | Uncorrected UMI (%) | 91.44 | 91.03 | 84.16 | 83.49 | 82.64 |
| | | Corrected UMI | 402 | 312 | 3,127 | 11,834 | 17,875 |
| | | Corrected UMI (%) | 0.03 | 0.02 | 0.06 | 0.11 | 0.14 |

| | | | | | |
|---|---|---|---|---|---|
| Total number of passed UMI | 1,465,706 | 1,175,994 | 4,417,663 | 8,754,892 | 10,949,369 |
| Total number of passed UMI (%) | 95.55 | 95.40 | 94.90 | 93.38 | 93.35 |

FLNC read count: number of Full-Length Non-Concatemer reads;
For cell barcode correction, “All” means CB have no QV filter, “CB QV>=0.95 (passed)” means have QV filter conditions before CB correction:
CB in allowlist: cell barcode can be found directly in the allowlists;
CB in allowlist (%): the percentage of “CB in allowlist” in FLNC read count;
Corrected CB: cell barcode has 1-Hamming distance with allowlists;
Corrected CB (%): the percentage of “Corrected CB” in FLNC read count;
Total number of corrected CB: sum of the “CB in allowlist” and “Corrected CB”;
Total number of corrected CB (%): the percentage of “Total number of corrected CB” in FLNC read count;
For UMI correction, “CB passed and UMI passed” means have QV filter conditions before UMI correction that CB QV must >=0.95 and have UMI filter conditions after correction that UMI must not “Discarded UMI”;
Discarded UMI: means UMI not passed basic quality filtering;
Discarded UMI (%): the percentage of “Discarded UMI” in FLNC read count;
Uncorrected UMI: means UMI have passed the basic quality filtering but no need correction;
Uncorrected UMI (%): the percentage of “Uncorrected UMI” in FLNC read count;
Corrected UMI (%): meet the UMI correction conditions and have correction;
Corrected UMI (%): the percentage of “Corrected UMI” in FLNC read count;
Total number of passed UMI: sum of the “Uncorrected UMI” and “Corrected UMI”;
Total number of passed UMI (%): the percentage of “Total number of passed UMI” in FLNC read count.

**Supplementary Table 3. The raw and filtered gene and UMI count of single cell in NGS, scISOseq, Linked-scISOseq, HIT-scISOseq gene matrix**

| | | NGS | | scISOseq | | Linked-scISOseq | HIT-scISOseq | |
|---|---|---|---|---|---|---|---|---|
| | Sample | s1 | s2 | s1 | s2 | s1 | s1 | s2 |
| Raw | Number of cells | 2171 | 2214 | 2171 | 2214 | 2171 | 2171 | 2214 |
| | Mean UMI count per cell | 14936 | 13740 | 413 | 300 | 1119 | 2138 | 2407 |
| | Median UMI count per cell | 12414 | 10024 | 370 | 232 | 932 | 1795 | 1779 |
| | Mean gene count per cell | 3127 | 2804 | 296 | 221 | 630 | 1042 | 1096 |
| | Median gene count per cell | 3189 | 2768 | 280 | 192 | 571 | 976 | 959 |
| | Total number of genes | 16154 | 16088 | 13946 | 13594 | 15161 | 16039 | 16310 |
| QC filter | Number of cells | 1723 | 1525 | 1658 | 1408 | 1735 | 1776 | 1599 |
| | Mean UMI count per cell | 15924 | 15466 | 514 | 438 | 1313 | 2451 | 3057 |
| | Median UMI count per cell | 14979 | 14683 | 467 | 390 | 1173 | 2208 | 2690 |
| | Mean gene count per Cell | 3485 | 3333 | 367 | 320 | 745 | 1208 | 1403 |
| | Median gene count per Cell | 3548 | 3525 | 342 | 294 | 691 | 1149 | 1341 |
| | Total number of genes | 13681 | 13588 | 10551 | 9928 | 12039 | 13084 | 13373 |

Number of cells: total number of cells in gene matrix;
Mean UMI count per cell: mean UMI detection of each cell in gene matrix;
Median UMI count per cell: median UMI detection of each cell in gene matrix;
Mean gene count per cell: mean type of isoform detection of each cell in gene matrix;
Median gene count per cell: median type of isoform detection of each cell in gene matrix;
Total gene of isoforms: sum of isoform type detection of all cell in gene matrix.

**Supplementary Table 4. The raw and filtered isoform and UMI count of single cell in scISOseq, Linked-scISOseq, HIT-scISOseq isoform matrix**

| | | scISOseq | | Linked-scISOseq | HIT-scISOseq | |
|---|---|---|---|---|---|---|
| | Sample | s1 | s2 | s1 | s1 | s2 |
| Raw | Number of cells | 2171 | 2214 | 2171 | 2171 | 2214 |
| | Mean UMI count per cell | 394 | 279 | 1317 | 2638 | 3006 |
| | Median UMI count per cell | 351 | 216 | 1094 | 2232 | 2267 |
| | Mean isoform count per cell | 296 | 209 | 874 | 1668 | 1837 |
| | Median isoform count per cell | 270 | 172 | 741 | 1438 | 1446 |
| | Total number of isoforms | 48786 | 37186 | 132705 | 236670 | 264228 |
| QC filter | Number of cells | 1598 | 1319 | 1702 | 1767 | 1533 |
| | Mean UMI count per cell | 459 | 382 | 1434 | 2769 | 3262 |
| | Median UMI count per cell | 416 | 346 | 1284 | 2513 | 3059 |
| | Mean isoform count per cell | 334 | 277 | 927 | 1714 | 1970 |
| | Median isoform count per cell | 310 | 254 | 840 | 1557 | 1851 |
| | Total number of isoforms | 24158 | 17839 | 70462 | 133064 | 150072 |

Number of cells: total number of cells in isoform matrix;
Mean UMI count per cell: mean UMI detection of each cell in isoform matrix;
Median UMI count per cell: median UMI detection of each cell in isoform matrix;
Mean isoform count per cell: mean type of isoform detection of each cell in isoform matrix;
Median isoform count per cell: median type of isoform detection of each cell in isoform matrix;
Total number of isoforms: sum of isoform type detection of all cell in isoform matrix.

**Supplementary Table 5. Shared cell barcode in each cell cluster between NGS and HIT-scISOseq data sets**

| Sample | Cell type | NGS cell counts | HIT-scISOseq cell counts | Shared cell barcode counts | Percentage of shared cell barcodes in NGS (%) | Percentage of shared cell barcodes in HIT-scISOseq (%) |
|---|---|---|---|---|---|---|
| s1 | Differentiated cells (k12+) | 440 | 462 | 438 | 99.55 | 94.81 |
| | Corneal basal cells (k14+) | 217 | 225 | 215 | 99.08 | 95.56 |
| | Limbal stem cells (k15+) | 624 | 634 | 619 | 99.20 | 97.63 |
| | Conjunctival cells (k4+) | 211 | 243 | 211 | 100.00 | 86.83 |
| s2 | Differentiated cells (k12+) | 392 | 421 | 392 | 100.00 | 93.11 |
| | Corneal basal cells (k14+) | 167 | 202 | 166 | 99.40 | 82.18 |
| | Limbal stem cells (k15+) | 506 | 545 | 501 | 99.01 | 91.93 |
| | Conjunctival cells (k4+) | 208 | 248 | 208 | 100.00 | 83.87 |

Shared cell barcodesdenote identical cells between NGS and HIT-scISOseq data sets that are within the same cell type and with the same cell barcode.
