## Supplementary Figures for "HIT-scISOseq: High-throughput and High-accuracy Single-cell Full-length Isoform Sequencing for Corneal Epithelium"

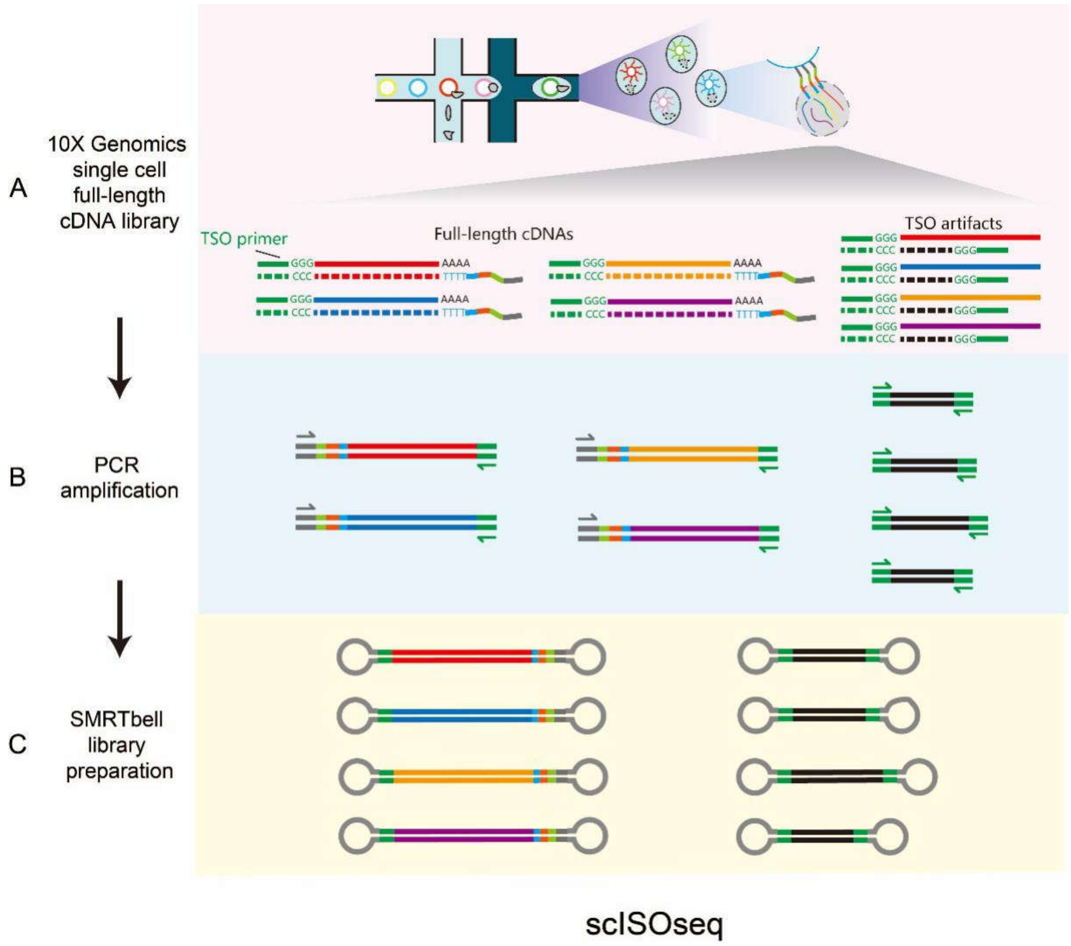


Figure S1. scISOseq and Linked-scISOseq workflow. (A) Overview of the 10X Genomics single cell full-length cDNA library construction. (B) cDNA amplification with a 3’ read 1 primer (grey) and TSO PCR primer (green). (C) SMRTbell library preparation.

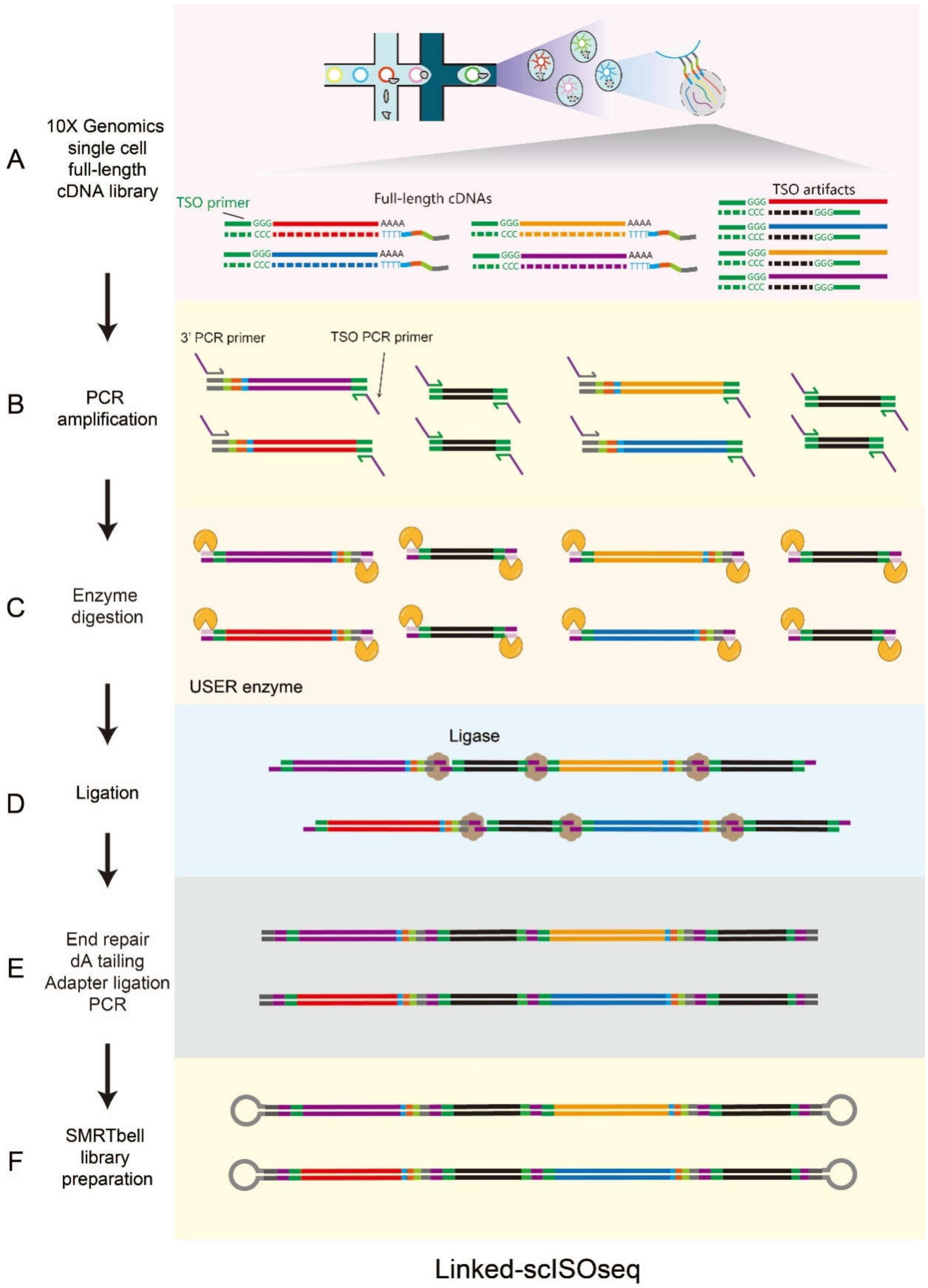


Figure S2. Linked-scISOseq workflow. (A) Overview of the 10X Genomics single cell full-length cDNA library construction. (B) cDNA amplification with a 3' PCR primer and TSO PCR primer. (C) Restriction digestion both ends primer to produce sticky end by USER enzyme. (D) and ligation of the cDNAs. (E-F) End repair, dA tailing, adapter ligation, and PCR enrichment of the ligated cDNAs , followed by SMRTbell library preparation.

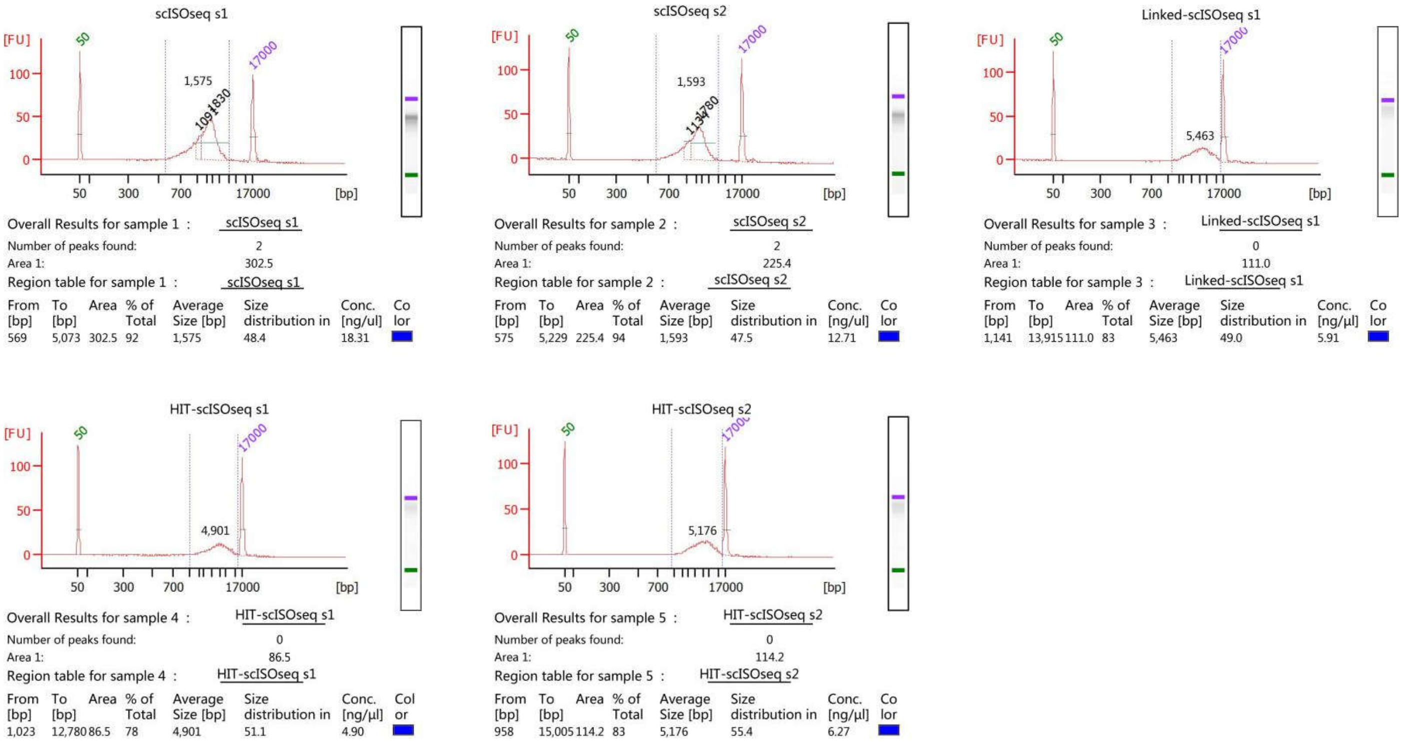


Figure S3. Agilent Bioanalyzer 2100 analysis showing results of the full-length cDNA libraries. These include scISOseq (A-B) cDNA library samples, Linked-scISOseq (C) library samples, and HIT-scISOseq (D-E) library samples QC and quantification.

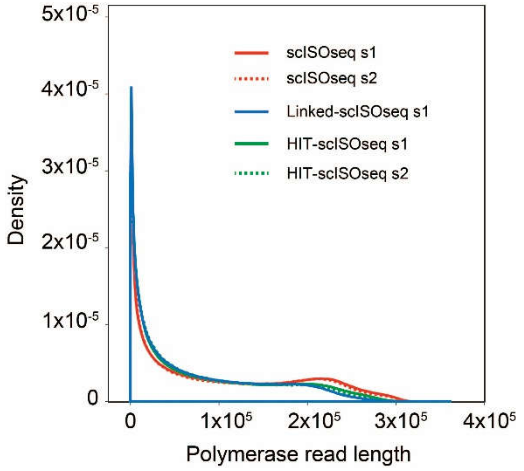


Figure S4. Polymerase read length distribution of scISOseq, Linked-scISOseq, and HIT-scISOseq cDNA library samples.

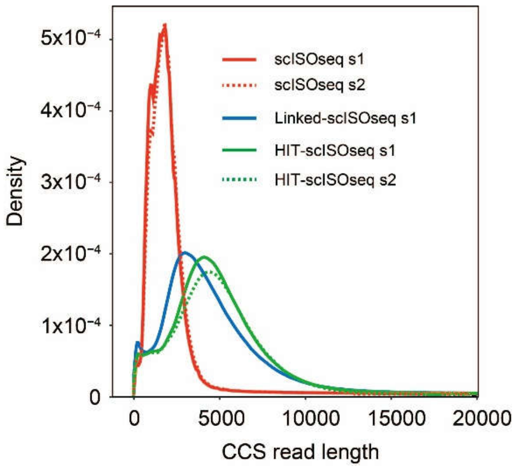


Figure S5. CCS read length distribution of scISOseq, Linked-scISOseq, and HIT-scISOseq cDNA library samples.

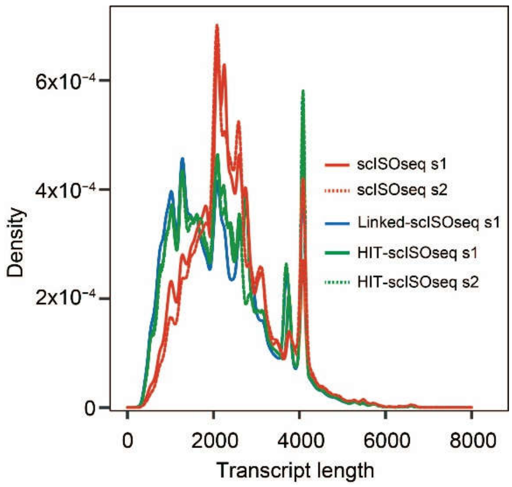


Figure S6. Collapsed reads (transcripts) length distribution of scISOseq, Linked-scISOseq, and HIT-scISOseq cDNA library samples.

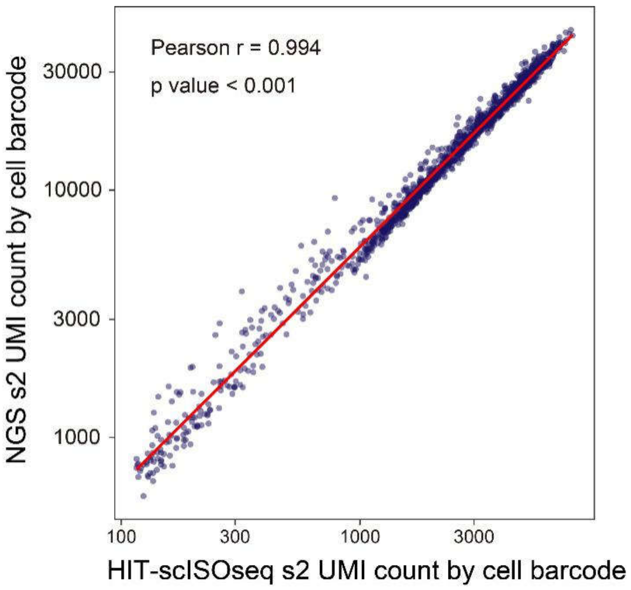


Figure S7. Correlation dot plot of replicate sample s2 of NGS (y axis) and HIT-scISOseq (x axis) UMI count by cell barcode. The correlation coefficient Pearson r, and P value are shown.

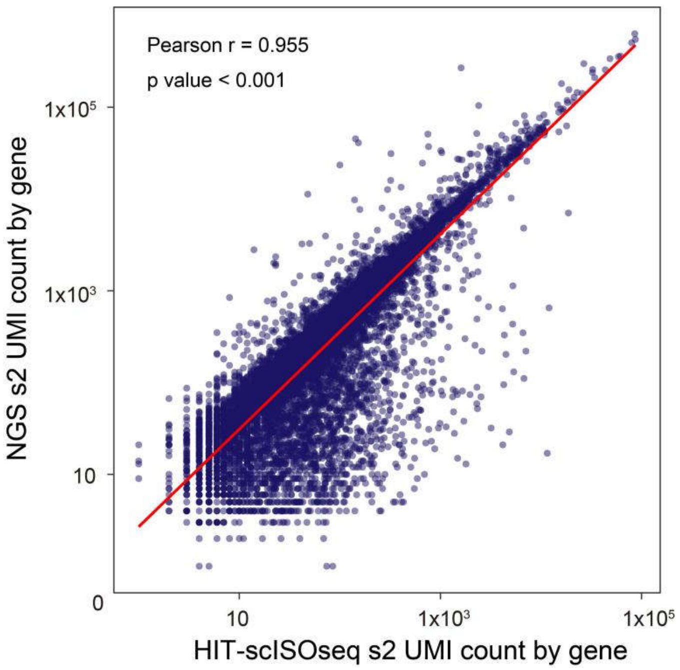


Figure S8. Correlation dot plot of replicate sample s2 of NGS (y axis) and HIT-scISOseq (x axis) UMI count by gene. The correlation coefficient Pearson r and P value are shown.

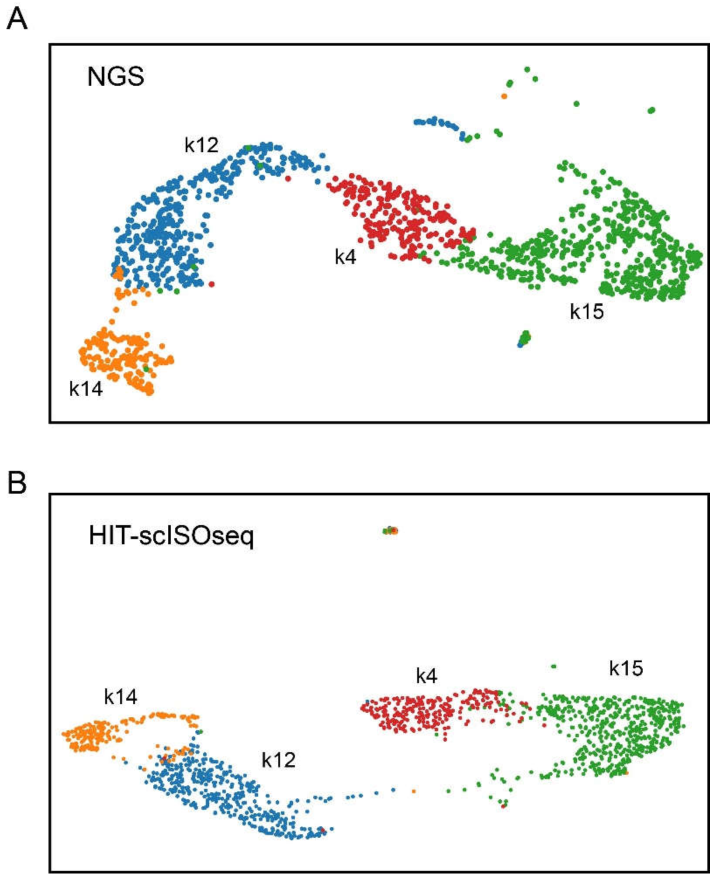


Figure S9. Clustering of single-cells from replicate sample s2 using NGS and HIT-scISOseq gene expression datasets. Both NGS (A) and HIT-scISOseq (B) data sets showing that the four main cell clusters could be successfully clustered (differentiated cells (K12+) = blue, corneal basal cells (K14+) = orange, limbal stem cells (K15+) = green, and conjunctival cells (K4+) = red).

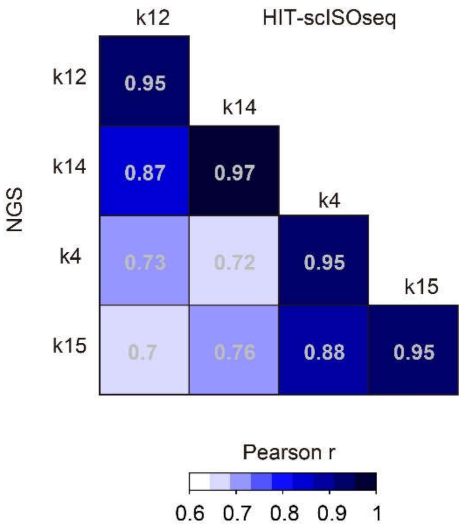


Figure S10. Gene expression correlation heatmap of each cell cluster between NGS and HIT-scISOseq data sets from replicate sample s2.

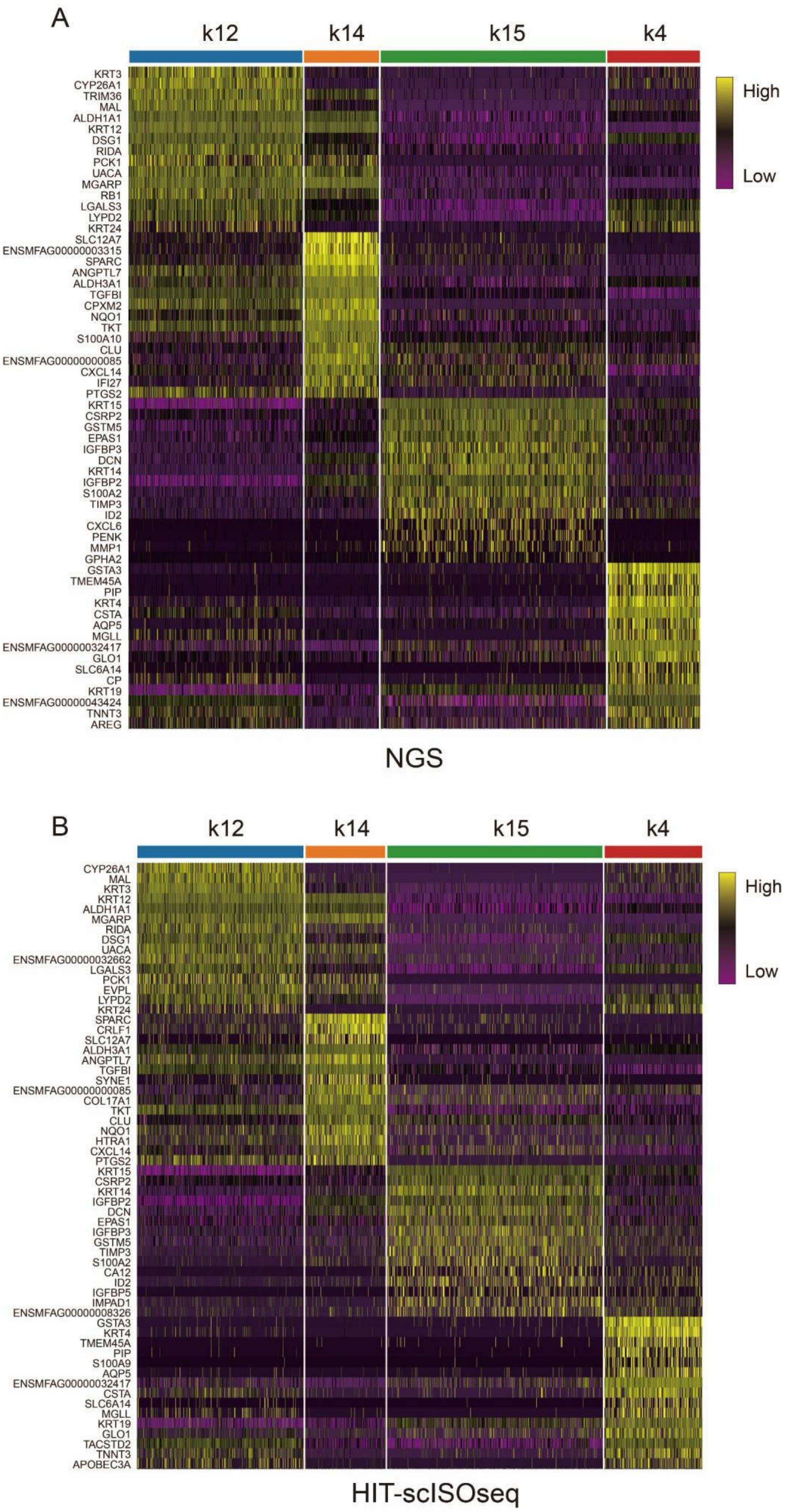


Figure S11. Marker gene expression heatmap of the four cell clusters. Data are generated from NGS (A) and HIT-scISOseq (B) data sets from replicate sample s2. Color gradient represents log-transformed and normalized counts scaled to a maximum of 1 per row. Upper bars represent the cell groups assignment for individual cells.

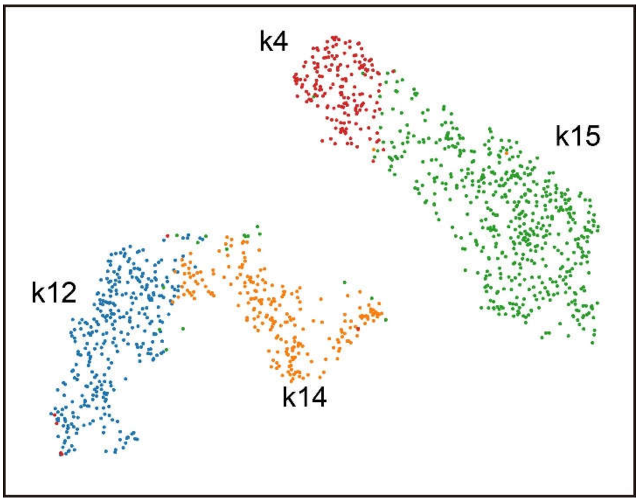


Figure S12. Cell type clustering based on HIT-scISOseq full-length isoform expression matrix in replicate sample s2. Four cell clusters are generated (differentiated cells (K12+) = blue, corneal basal cells (K14+) = orange, limbal stem cells (K15+) = green, and conjunctival cells (K4+) = red).

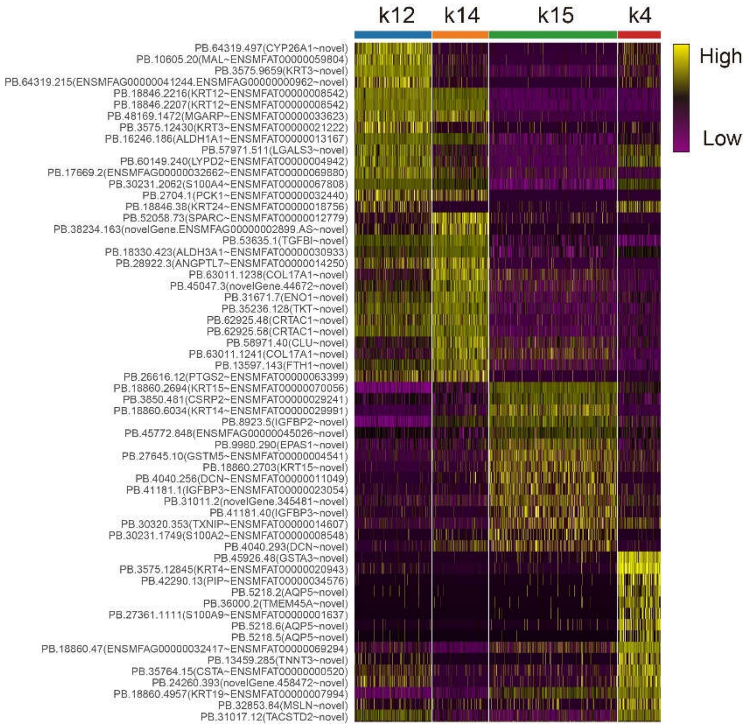


Figure S13. Single cell maker isoforms expression heatmap of HIT-scISOseq data from replicate sample s2. Color gradient represents log-transformed and normalized counts scaled to a maximum of 1 per row. Upper bars represent cell cluster' assignment for individual cells.

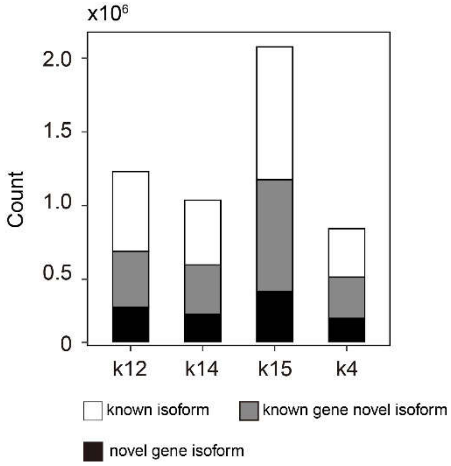


Figure S14. Distribution of known and novel isoforms in each cell cluster of HIT-scISOseq data from replicate sample s2. Isoform classification are color coded as follows: known isoforms (white), known gene's novel isoforms (grey) and novel gene's isoforms (black).

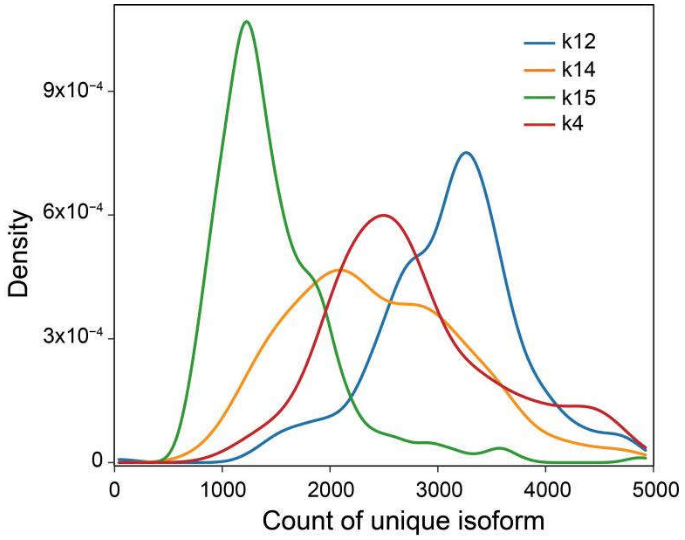


Figure S15. Distribution of expressed unique isoform numbers (isoforms with minimal UMI count of 3) of individual cells in each cell clusters of HIT-scISOseq data from replicate sample s2 (differentiated cells (K12+) = blue, corneal basal cells (K14+) = orange, limbal stem cells (K15+) = green, and conjunctival cells (K4+) = red). Y-axis for density of cell barcode count.

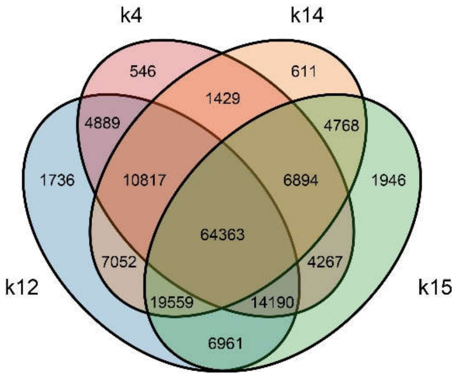

Figure S16. Venn plot of cell-type-specific isoforms of HIT-scISOseq data from replicate sample s2.
